# In situ modeling of acquired resistance to RTK/RAS pathway targeted therapies

**DOI:** 10.1101/2023.01.27.525958

**Authors:** Nancy E. Sealover, Patricia T. Theard, Amanda J. Linke, Jacob M. Hughes, Brianna R. Daley, Robert L. Kortum

**Affiliations:** Department of Pharmacology and Molecular Therapeutics, Uniformed Services University of the Health Sciences, Bethesda, MD

**Keywords:** acquired resistance, EGFR, RAS, MEK, G12C, SHP2, RTK

## Abstract

Intrinsic and acquired resistance limit the window of effectiveness for oncogene-targeted cancer therapies. Preclinical studies that identify synergistic combinations enhance therapeutic efficacy to target intrinsic resistance, however, methods to study acquired resistance in cell culture are lacking. Here, we describe an *in situ* resistance assay (ISRA), performed in a 96-well culture format, that models acquired resistance to RTK/RAS pathway targeted therapies. Using osimertinib resistance in *EGFR*-mutated lung adenocarcinoma (LUAD) as a model system, we show acquired resistance can be reliably modeled across cell lines with objectively defined osimertinib doses. We further show that acquired osimertinib resistance can be significantly delayed by inhibition of proximal RTK signaling using two distinct SHP2 inhibitors. Similar to patient populations, isolated osimertinib-resistant populations showed resistance via enhanced activation of multiple parallel RTKs so that individual RTK inhibitors did not re-sensitize cells to osimertinib. In contrast, inhibition of proximal RTK signaling using the SHP2 inhibitor RMC-4550 both re-sensitized resistant populations to osimertinib. Similar, objectively defined drug doses were used to model resistance to additional RTK/RAS pathway targeted therapies including the KRAS^G12C^ inhibitors adagrasib and sotorasib, the MEK inhibitor trametinib, and the farnesyl transferase inhibitor tipifarnib. These studies highlight the tractability of in situ resistance assays to model acquired resistance to targeted therapies and provide a framework for assessing the extent to which synergistic drug combinations can target acquired drug resistance.

**Highlights:**

- Acquired resistance to RTK/RAS pathway members can be modeled in situ
- SHP2 inhibitors reduce the development of acquired osimertinib resistance
- Isolated osimertinib-resistant populations show hyperactivation of multiple RTKs
- SHP2 inhibitors re-sensitize resistant populations to osimertinib treatment

## Introduction

Lung cancer is the leading cause of cancer-related death worldwide; adenocarcinomas are the most common subtype of lung cancer^1^. Oncogenic driver mutations in the RTK/RAS/RAF pathway occur in 75-90% of lung adenocarcinomas (LUAD). With enhanced understanding of the oncogenic driver mutations causing disease, oncogene-targeted therapies have substantially improved patient outcomes not only for patients with LUAD, but across the spectrum of cancer types. However, in most cases resistance to targeted therapeutics develops necessitating novel approaches that can either delay therapeutic resistance or treat resistant cancers.

Activating mutations in the epidermal growth factor receptor (EGFR) drive oncogenesis in 15-30% of LUADs^1^. For patients with EGFR-mutated tumors, the EGFR-TKI osimertinib markedly enhances progression free and overall survival compared to standard chemotherapy^2, 3^, however, both intrinsic and acquired osimertinib resistance limit patient survival. Acquired osimertinib resistance is most often driven by RTK/RAS pathway reactivation^4^, either via secondary EGFR mutations or enhanced signaling through multiple parallel RTKs including MET, AXL, FGFR, HER2/3, and IGF1-R^4–13^. This heterogeneity of RTKs that can drive osimertinib resistance, along with the finding that at rapid autopsy patient tumors often harbor multiple modes of RTK-dependent osimertinib resistance^14^, may limit the up-front effectiveness of combining osimertinib with a second RTK inhibitor.

Upregulation of heterogeneous RTK activity has been documented as an intrinsic and acquired mechanism of resistance in response to mono-therapeutics designed to target RTK/RAS driven cancers. The non-receptor protein tyrosine phosphatase SHP2 has been suggested to support RTK activity in drug resistant cell populations as a ubiquitous regulator of proximal RTK signaling. While combining SHP2 inhibition with a RTK or RAS inhibitor has been shown to be effective, use of SHP2 inhibitors to block intrinsic or acquired resistance is lacking in models with directly measured heterogeneous RTK upregulation. Further, many of these publications do not assess the effect of SHP2 inhibition on a broad time scale representative of acquired resistance. We therefore aimed to develop a model to study the role of SHP2 inhibition in cells that had developed resistance to RTK targeted therapies over 6-12 weeks and capture the effect of SHP2 inhibition in cells that have heterogeneous RTK upregulation as a response to previous oncogene inhibition.

Historically, combination therapies have been based on identifying two or more active single agents with distinct mechanisms of action^15^, and most current combination therapies include an oncogene-targeted therapy and a second drug that targets an independent mechanism that, if left targeted, causes to resistance to the first agent^16, 17^. While many of these approaches succeed in increasing windows of progression free survival (PFS) in patients, resistance still emerges, often due to the strikingly heterogeneous mechanisms of acquired drug resistance both intratumorally^14, 18^ and across patients^19, 20^. Thus, novel approaches are needed to identify combination therapies that not only address a single mechanism of acquired resistance but simultaneously prevent most common mechanisms of resistance to a given targeted therapy. This need has led to a shift toward groups attempting to identify synergistic drug combinations to treat cancers with specific genetic profiles^15^.

Preclinical studies designed to identify synergistic drug combinations are excellent at capturing the success of a drug treatment within a timeframe of 1-14 days. Unfortunately, few studies extend beyond assessing this initial window of efficacy to determine whether combinations can prevent the development of acquired resistance. For example, RNAi and CRISPR screens have been widely successful in identifying novel and often unexpected partners for combination therapies^21–23^, however, the timeframe of these approaches biases them toward identifying and assessing secondary drug targets that will limit intrinsic, but not acquired, resistance. Further, those studies that do assess acquired resistance *in situ* are limited to the assessment of a few cell lines established by dose-escalation over multiple months (reviewed in^24^) (e.g.^25–29^) rather than determining the extent to which inhibiting a secondary drug target can delay the onset of resistance. Therefore, a framework of pre-clinical experiments to assess initial efficacy combined with extended *in vitro* approaches such as 6-16 week proliferation outgrowth assays^30, 31^ and time-to-progression (TTP) assays^23^ can provide robust evidence for proposing effective and rational combination therapies that is accessible and scalable.

Here, we describe an *in situ* resistance assay (ISRA) that allows us to (i) assess the development of acquired resistance in a large cohort of individual cultures, (ii) isolate multiple therapy-resistant polyclonal cell populations for biochemical analysis, and (iii) test the effectiveness of combination therapies to delay the development of acquired resistance. We used *EGFR*-mutated LUAD cell lines as a model to assess the timeframe of acquired resistance to the third generation tyrosine kinase inhibitor (TKI) osimertinib in an *in situ* resistance assay (ISRA) that combines elements from proliferation outgrowth ^31^ and TTP ^23^ assays, and allows for screening of drug combinations in large numbers of individual cell populations using a 96-well format. We show that cells isolated from osimertinib ISRAs remain osimertinib resistant and have developed acquired resistance via upregulation of multiple parallel RTKs. Using a panel of inhibitors targeting multiple receptor tyrosine kinases (RTKs) and RTK-associated proteins, we found that inhibiting proximal RTK signaling using the SHP2 inhibitor RMC-4550 ^32^ most consistently increased both efficacy and potency of osimertinib across all osimertinib resistant cell lines. In addition to re re-sensitizing osimertinib resistant lines to osimertinib, we further showed that SHP2 inhibition both delayed the onset and reduced the overall frequency of osimertinib resistant cultures. This model supports the suggested mechanism that SHP2 acts a central signaling node for RTK upregulation during acquired drug resistance. Finally, we demonstrated the broad applicability of ISRAs to model acquired resistance to multiple targeted including the KRAS^G12C^ inhibitors adagrasib and sotorasib, the MEK inhibitor trametinib, and the farnesyl transferase inhibitor tipifarnib in multiple cell line models.

## Results

### Development of an assay to model acquired resistance in situ

We sought to develop an *in situ* resistance assay (ISRA) that modeled the development of acquired resistance to targeted therapies and could be performed in a multi-well setup to mimic a multiple-subject trial testing the effectiveness of drug combinations to limit the development of resistance. To develop and optimize the ISRA, we used a panel of *EGFR*-mutated lung adenocarcinoma (LUAD) lines treated with the third-generation TKI osimertinib, as the mechanisms driving osimertinib resistance in LUAD are well established^4–13^. To establish the doses of osimertinib that inhibits survival (≥ EC_50_) in the panel of *EGFR*-mutated cell lines, H1975, H827, PC9, and PC9-TM (PC9 cells with an acquired T790M mutation) cells were treated with increasing doses of osimertinib and cell viability was assessed four days after treatment (Fig. 1A and B). The doses of osimertinib that inhibited survival were similar across all *EGFR*-mutated cell lines [EC_50_ (17-46 nM) - EC_85_ (200-450 nM)], allowing us to use similar doses in each cell line to determine the concentration of osimertinib that caused prolonged growth arrest (> 6 weeks) prior to cell outgrowth in a majority of cell populations for each cell line. Cells were plated at low density (250 cells/well; <10% confluent) in multiple 96-well plates and each plate was either left untreated (to assess the time required for normal cell outgrowth) or treated with a single inhibitory dose of osimertinib (EC_50_-EC_85_ range; 30, 50, 150, or 300 nM) for up to 12 weeks. The inner 60 wells of each plate were visually assessed weekly and wells that were ≥ 50% confluent were scored as resistant to that dose of osimertinib. Lower doses of osimertinib corresponding to the EC_50_-EC_75_ caused a 1–3-week delay in the outgrowth of cells, whereas cells treated with ≥ EC_80_ of osimertinib showed prolonged growth arrest before regaining proliferative capacity (Fig. 1C) under continuous osimertinib treatment, indicative of cells that had become osimertinib resistant. These results suggest that osimertinib resistance can be modeled *in situ* using doses of osimertinib with ≥ EC_80_ for growth inhibition in *EGFR*-mutated LUAD cell lines.

**Fig 1.**
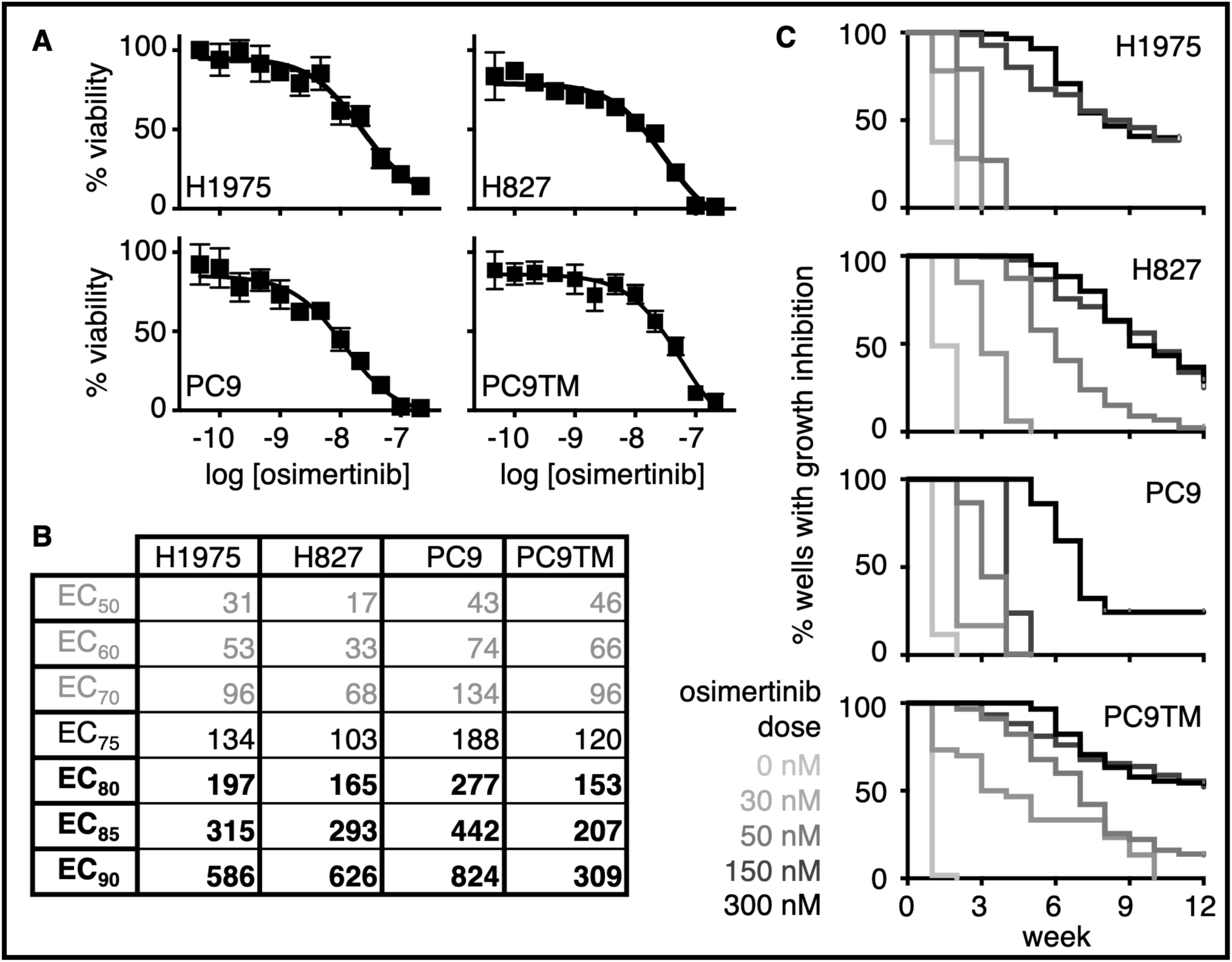
Modeling osimertinib resistance in situ. **A-B**. Dose response curves (A) and EC_50_ – EC_90_ values (B) for the indicated *EGFR*-mutated LUAD cell lines treated with osimertinib under anchorage-dependent conditions. Data in A are mean ± sd from three independent experiments. **C**. EGFR-mutated cell lines were plated at low density (250 cells/well) in replicate 96-well plates, and each plate was treated with the indicated dose of osimertinib. Wells were fed and assessed weekly for outgrowth, wells that were > 50% confluent were scored as resistant to the given dose of osimertinib. Data are plotted as a Kaplan-Meyer survival curve. Plates treated with ≥ the EC_80_ osimertinib dose (see **bold values** in B) showed an extended inhibition of growth followed by outgrowth of individual colonies consistent with true drug resistance. Data in C are combined from three independent trials.

Acquired osimertinib resistance is commonly driven by RTK/RAS pathway reactivation^4^ either via secondary EGFR mutations or enhanced signaling through one or more parallel RTKs including MET, AXL, FGFR, HER2/3, and IGF1-R. Given the multiplicity of RTKs capable of driving osimertinib resistance, we posited that inhibition of proximal RTK signaling, via SHP2 inhibition, would limit RTK-dependent osimertinib resistance and thereby prolong the initial window of osimertinib therapy. To directly assess the extent to which SHP2 inhibition could limit the development of acquired osimertinib resistance, we performed ISRAs in *EGFR*-mutated H1975 and PC9 cells treated with osimertinib either alone in combination with two distinct SHP2 inhibitors [RMC-4550 or SHP099 (Fig. 2)]. When treated with osimertinib alone at either 150 or 300 nM, H1975 cells show a median time to outgrowth of 9 weeks with 63% of populations eventually becoming osimertinib. Combining RMC-4550 or SHP099 with osimertinib significantly inhibited the development of osimertinib resistance, with only 3% (RMC-4550) or 21% (SHP099) of populations treated with combination therapy showing outgrowth at 12-weeks.

**Fig 2.**
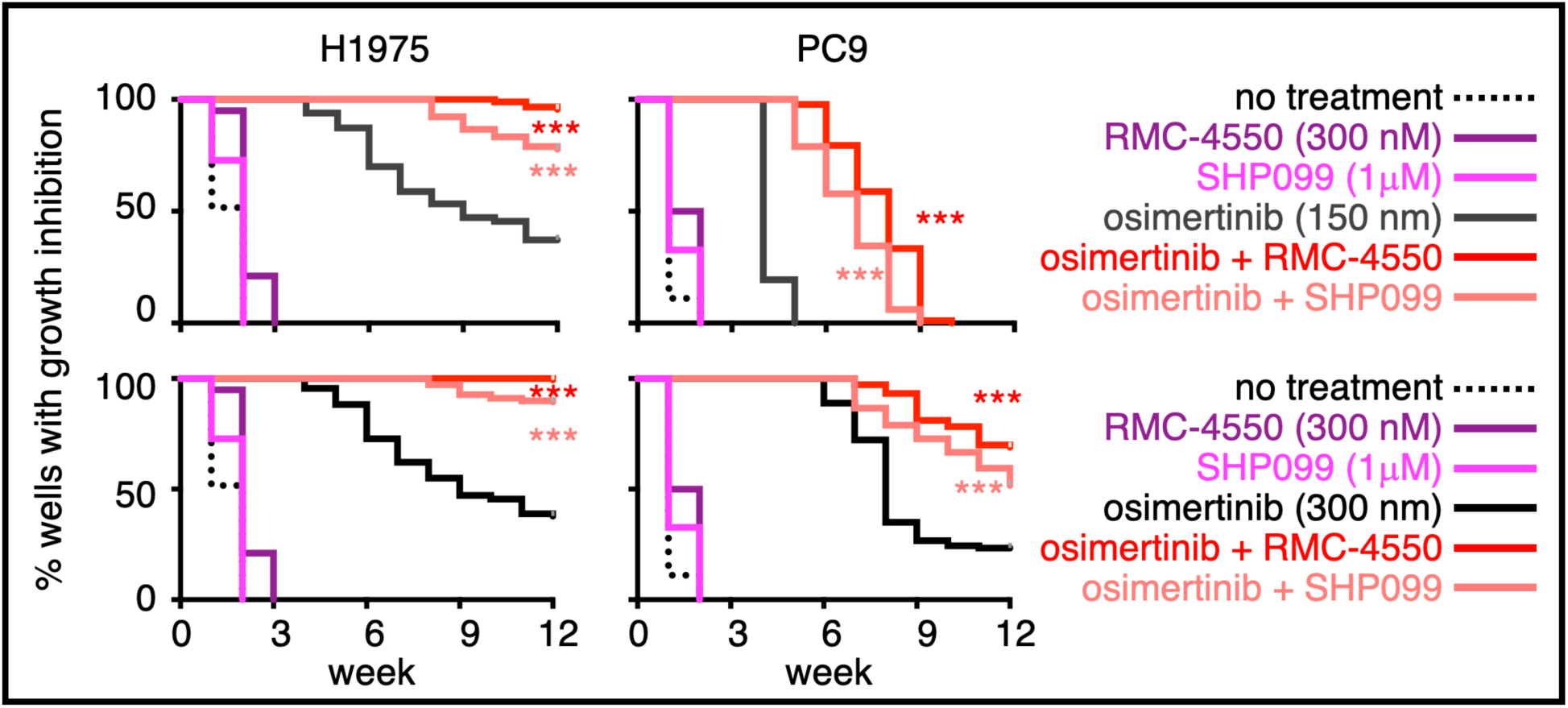
SHP2 inhibition limits the development of osimertinib resistance. *EGFR*-mutated H1975 and PC9 cells were plated at low density (250 cells/well) in replicate 96-well plates, and each plate was either left untreated (black, dashed) or treated with osimertinib alone at 150 (dark grey) or 300 nM (black), a SHP2 inhibitor alone (RMC-4550 at 300 nM or SHP099 at 1 mM) (purple), or the combination of osimertinib + RMC-4550 or SHP099 (red). Wells were fed and assessed weekly for outgrowth, wells that were > 50% confluent were scored as resistant to the given dose of osimertinib-resistant. Data are plotted as a Kaplan-Meyer survival curve. *** p<0.001 vs single drug treatment. Data are combined from three independent trials.

Similar results were observed in PC9 cells treated with 300 nM osimertinib, cells treated with osimertinib alone showed at median time to outgrowth of 8 weeks with 76% of total populations becoming osimertinib resistant. Combining RMC-4550 or SHP099 with 300 nM osimertinib significantly inhibited the development of osimertinib resistance, with only 30% (RMC-4550) or 41% (SHP099) of populations treated with combination therapy showing outgrowth at 12-weeks. Further, while PC9 cells treated with 150 nM osimertinib show rapid outgrowth (median time to outgrowth of 4 weeks), this outgrowth was significantly delayed by either RMC-4550 or SHP099. Overall, these data show both the utility of ISRAs to test the exent to which therapeutic combinations can inhibit the development acquired resistance and show that SHP2 inhibition may significantly prolong the window for osimertinib therapy.

### RTK phosphorylation is heterogeneously upregulated in osimertinib-resistant cells

We next wanted to assess if the mechanisms driving acquired osimertinib resistance in cells isolated from IRSAs would be predictive of sensitivity to SHP2 inhibition. We isolated and expanded a cohort of osimertinib-resistant populations (OR1-8) from H1975 individual wells treated with 150 nM osimertinib to investigate the extent to which (i) individual populations maintained osimertinib resistance and (ii) the mechanisms driving osimertinib resistance in cells isolated from continued drug treatment were similar to those seen in patient populations and predictive of sensitivity to proximal RTK pathway inhibition using a SHP2 inhibitor. Dose-response studies showed that osimertinib was much less potent in all eight OR populations compared to H1975 parental cells (Fig. 3A and fig. S1A), with >40% of cells surviving when treated with 1M osimertinib compared to < 20% of parental controls. Further, in ISRA assays using 150 nM osimertinib, all eight populations showed osimertinib resistance (Fig. 3B and fig. S1B). While it took 9-weeks for 50% of parental H1975 wells to become osimertinib resistant and only 70% became resistant in the 12-week assay, OR1-8 populations showed outgrowth in 100% of wells within 3-5 weeks (Fig. 3B and fig. S1B). These data confirm that cell populations isolated from ISRAs maintain drug resistance.

**Fig 3.**
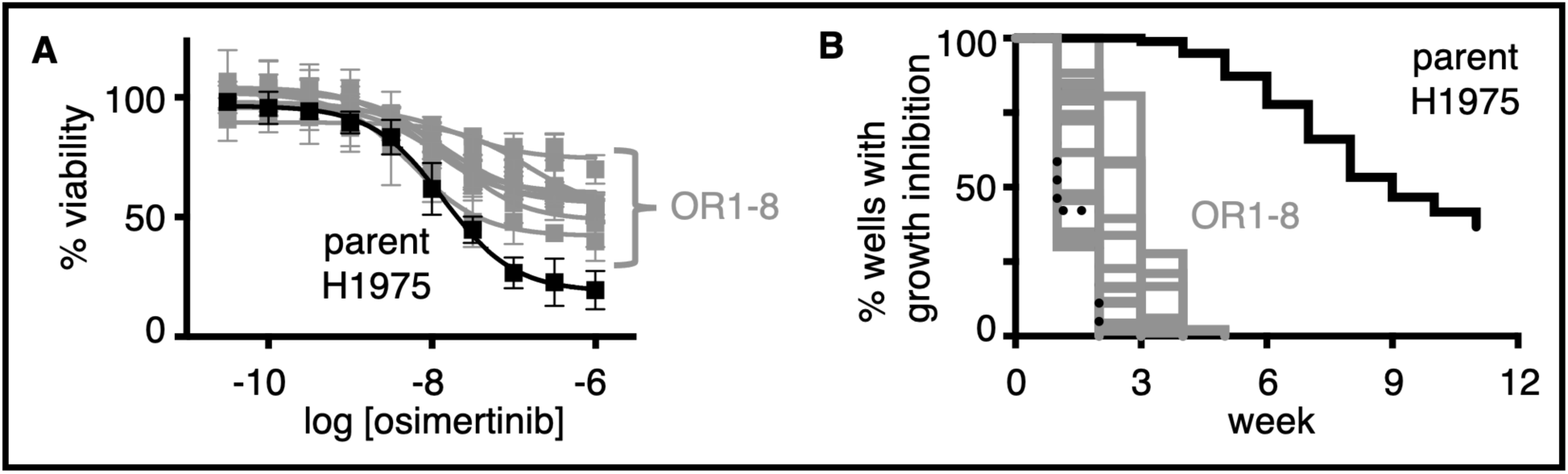
Populations isolated from in situ resistance assays maintain long-term osimertinib resistance. **A-B.** Dose response curves assessing osimertinib sensitivity at 96-h (A) and in situ resistance assays (B) in parental H1975 cells (black) and osimertinib-resistant H1975 populations OR1-OR8 (grey) isolated from wells that became confluent after treatment with 150 nM osimertinib for ≥ six weeks. Curves comparing individual OR lines to the parental H1975 population are shown in Fig. S1. Data in A are mean ± sd from three independent experiments. Data in B are combined from three independent trials.

To assess whether, similar to patient samples, OR populations isolated from ISRAs showed hyperactivation of parallel RTKs, we evaluated OR1-8 populations for changes in magnitude of receptor tyrosine phosphorylation compared to parental H1975 cells using phospho-tyrosine arrays (Fig. 4A and fig. S2A). OR1-8 populations showed increased phosphorylation of multiple RTKs; AXL and FGFR2-α were most consistently phosphorylated across OR1-8 populations with additional RTKs including c-MET, RET, IGF-1R, c-KIT, and TIE2 and non-receptor kinases ABL and SRC family kinases (SFKs) showing increased phosphorylation in at least two OR populations (Fig. 4A and fig. S2A). These data show that osimertinib-resistant populations isolated from ISRAs show reactivation of multiple RTKs, similar to the multiple mechanisms of RTK reactivation observed in osimertinib-resistant tumors from rapid autopsy studies^14^.

**Fig 4.**
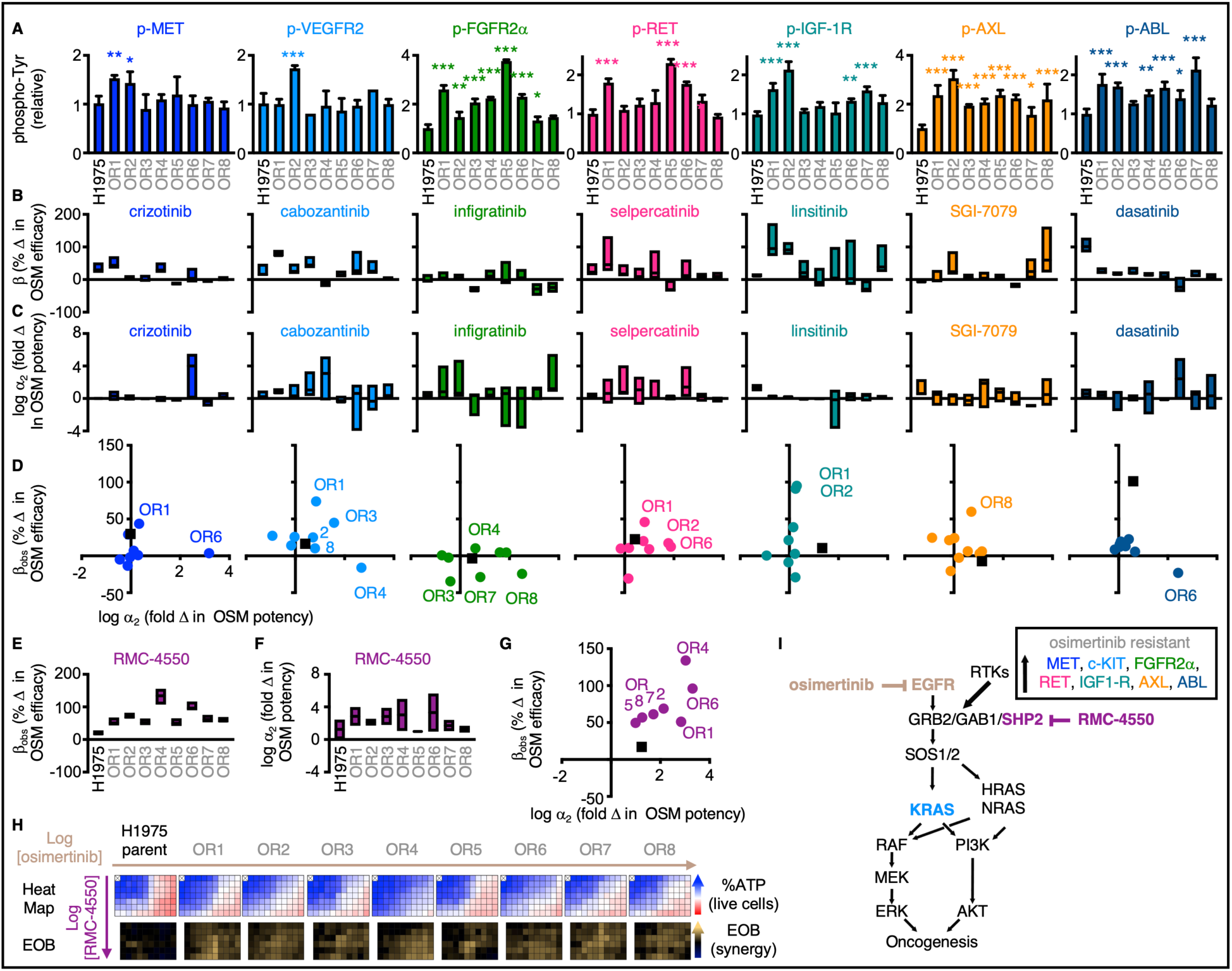
Osimertinib resistant populations show hyperactivation of multiple RTKs and are sensitive to SHP2 inhibition. **A.** Quantitation of tyrosine phosphorylation of the indicated RTKs in whole cell lysates from osimertinib-resistant H1975 populations OR1-8 relative to parental controls, samples were assessed for RTK phosphorylation using a Human RTK phosphorylation antibody array (RayBiotech, complete array data in Fig. S2). Data are mean ± sd from three replicate wells of the RTK array. **B-G.** Quantitation of the change in % osimertinib efficacy (**B, E**, ý_obs_), fold change in osimertinib potency (**C, F**, log α_2_), and a plot in the change in osimertinib potency (log α_2_) versus efficacy (ý_obs_) **(D, G)** from dose-response experiments assessing combined treatment of parental H1975 cells or OR1-8 populations with osimertinib +/- the indicated second RTK or RTK/RAS pathway inhibitor. For **D**, cells in the upper right quadrant (positive b_obs_ and log a_2_) represent the most promising osimertinib combination for a given cell population. Data are presented as the **H.** Parental and osimertinib-resistant H1975 populations were treated with a matrix of doses of osimertinib +/- the SHP2 inhibitor RMC-4550 for four days and cell viability was assessed using CellTitre glo. Excess over Bliss was calculated for each dose combination as a measure of synergy. **I.** Schematic showing hyperactivation of multiple RTKs driving acquired osimertinib resistance. Inhibition of proximal RTK signaling using a SHP2 inhibitor, but not individual RTK inhibitors, synergizes with osimertinib by enhancing osimertinib efficacy and potency.

### SHP2 inhibition re-sensitizes osimertinib resistant cells to osimertinib regardless of specific receptor activity

The current treatment paradigm for patients with osimertinib-resistant tumors is to determine secondary therapy based on the presence of druggable mechanism(s) of osimertinib resistance, most often using inhibitors of mutated or amplified RTKs. To determine the extent to which treatment with inhibitors of RTKs or non-receptor kinases showing hyperactivation in osimertinib resistant cells re-sensitized these cells to osimertinib, we used a panel of RTK inhibitors and the ABL/SFK inhibitor dasatinib; primary and secondary targets of each inhibitor are shown in Fig. S2B. We treated OR1-8 populations with either with increasing doses of osimertinib +/- 100 or 300 nM a second inhibitor or increasing doses of the second inhibitor +/- 10 nM osimertinib (Fig. S3). Synergistic drug-drug interactions were assessed by Multi-dimensional Synergy of Combinations (MuSyC) to deconvolute synergistic potency [Log(α_2_)] and efficacy (ý_obs_) _33,34_; optimal drug combinations show positive values for both Log(α_2_) and ý_obs_ and fall in the upper right quadrant of plots for these two indices of synergy. While individual combinations showed modest increases in syenrgistic potency and efficacy in specific cell lines (crizotinib and increased11ib in OR1; linsitinib in OR1 and OR2; SGI-7079 in OR8), overall the effects of specific RTK inhibitors or dasatinib was heterogenous among OR cell lines and not correlated with increased RTK signaling of the major target of any of the inhibitors tested (Fig. 4B, 4C, and 4D). In contrast, inhibition of proximal RTK signaling using the SHP2 inhibitor RMC-4550 increased the potency [Log(α _2_)] and efficacy (ý_obs_) of osimertinib in all OR1-8 populations (Fig. 4E-G). To further characterize the extent to which RMC-4550 synergized with osimertinib in osimertinib-resistant populations, we treated parental H1975 or OR1-8 with increasing doses of RMC-4550 and/or osimertinib in a 6×10 matrix of drug combinations and assessed for synergistic killing after 96 hours treatment by Bliss Independence (Fig. 4H). Here again, SHP2 inhibition markedly enhanced the killing effects of osimertinib in OR1-8 cells, showing an increased excess over Bliss across the matrix of drug combinations (Fig. 4H). These data suggest that inhibiting proximal RTK signaling has the potential to overcome RTK-driven osimertinib resistance in *EGFR*-mutated cancers and demonstrates a therapeutic role for inhibitors of proximal RTK signaling, including SHP2 inhibitors, in tumros with reported heterogeneous RTK upregulation in response to prior therapy (Fig. 4I).

### Modeling resistance to KRAS^G12C^ inhibitors, trametinib, and tipifarnib

Oncogenic driver mutations in the RTK/RAS/RAF pathway occur in 75-90% of LUAD. Similar to *EGFR*-mutated LUADs treated with osimertinib, acquired resistance to targeted therapies limits the overall window of their effectiveness to treat patients with LUAD. We sought to determine the extent to which ISRAs could be used as a general model of acquired resistance to agents targeting the RTK/RAS pathway. We therefore sought to model acquired resistance to the covalent KRAS^G12C^ inhibitors sotorasib (Fig. 5A) and adagrasib (Fig. 5B) in *KRAS^G12C^*-mutated (H358, H1373) LUAD cells, the MEK inhibitor trametinib in both *KRAS (non-G12C)*-mutated (H727) and *NF1*-LOF (H1838) LUAD cells (Fig. 5C), the farnesyltransferase inhibitor (FTI) tipifarnib in *HRAS*-mutated LUAD (H1915) and rhabdomyosarcoma (SMS-CTR) cells (Fig. 5D). To establish the doses of each inhibitor that inhibits survival (≥ EC_50_) in the panel of RTK/RAS*-*pathway mutated cell lines, cells were treated with increasing doses of the indicated inhibitor and cell viability was assesses four days after treatment (Fig. S4); for KRAS^G12C^ and MEK inhibitors dose response curves were performed in 3D culture as *KRAS*-mutated cells show variable responses to KRAS expression and KRAS^G12C^ inhibition in 2D culture^35–44^ (EC_50_ – EC_90_ values for each drug/cell line combination are shown in Fig. 5E). ISRAs were then performed using inhibitory doses (∼EC_50_ – EC_85_) for each inhibitor/cell line combination. For each cell line, lower doses of the targeted inhibitor corresponding to the EC_50_-EC_75_ caused a 1–3-week delay in the outgrowth of cells, whereas cells treated with ≥ EC_80_ of inhibitor showed prolonged growth arrest before regaining proliferative capacity (Fig. 5A-D) indicating that these wells had become resistant to the indicated therapeutic. These data show that, in addition to modeling osimertinib, ISRAs can be used to model acquired resistance to multiple therapies targeting the RTK/RAS/ERK pathway.

**Fig 5.**
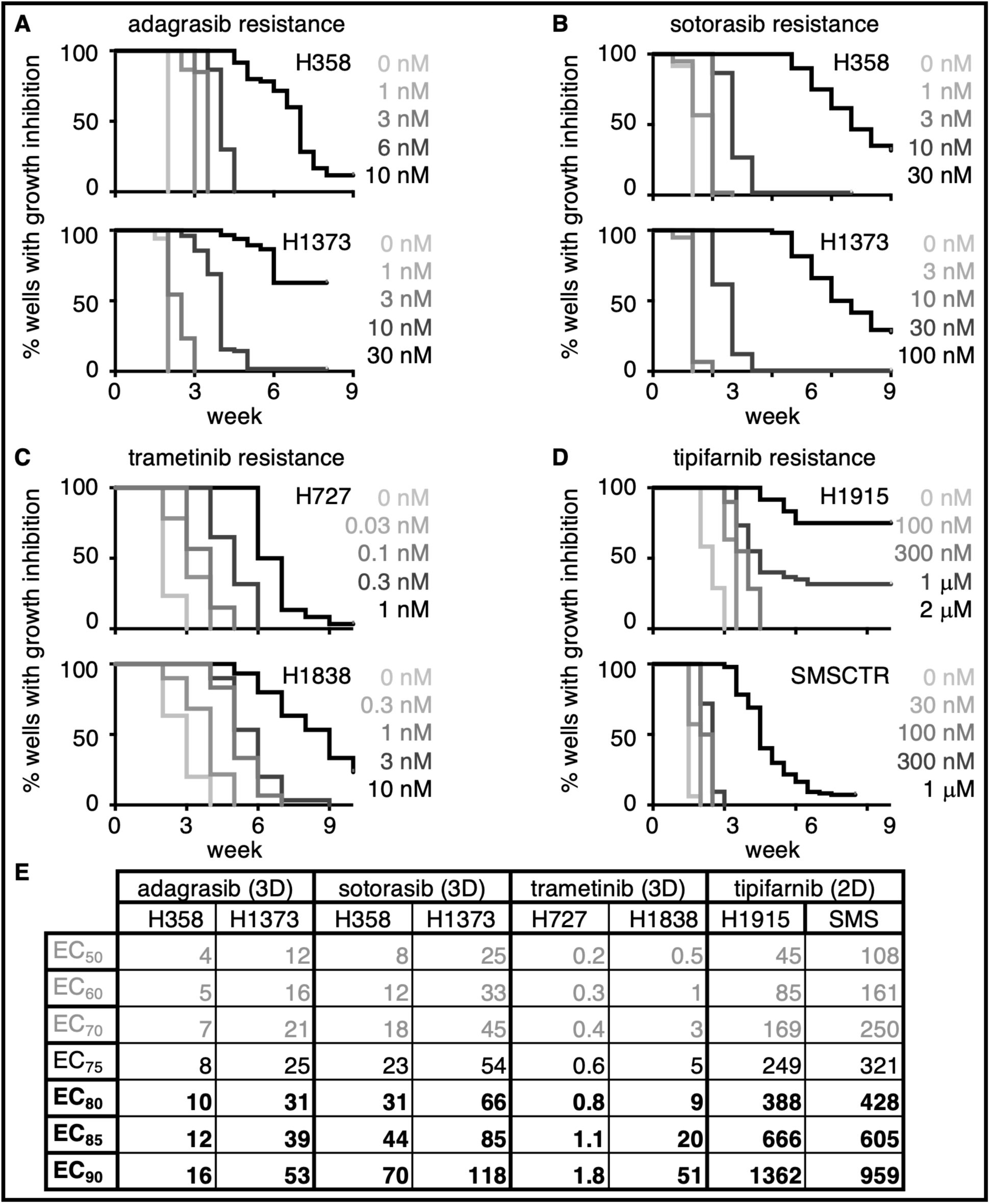
The in situ resistance assay can be used to model resistance to RTK/RAS pathway targeted therapies. **A-D**. *KRAS^G12C^*-mutated H358 and H1373 cells (A, B), *KRAS^G12C^*-mutated H727 and NF1-LOF H1838 cells (C), or HRAS-mutated H1915 and SMSCTR cells were plated at low density (250 cells/well) in replicate 96-well plates, and each plate was either left untreated or treated with the indicated dose of sotorasib (A), adagrasib (B), trametinib (C), or tipifarnib (D). Wells were fed and assessed weekly for outgrowth, wells that were > 50% confluent were scored as resistant to the given dose of osimertinib-resistant. Data are plotted as a Kaplan-Meyer survival curve. Plates treated with ≥ the EC_80_ dose of a given targeted inhibitor (see bold values in E) showed an extended inhibition of growth followed by outgrowth of individual colonies consistent with true drug resistance. **E.** EC_50_ – EC_90_ values from dose response curves assessing sotorasib (H358, H1373), adagrasib (H358, H1373), trametinib (H727, H1838), or tipifarnib (H1915, SMSCTR) sensitivity. Sotorasib, adagrasib, and trametinib sensitivities were assessed in 3D cultured spheroids; tipifarnib sensitivity was assessed in 2D adherent culture.

## Discussion

Oncogene-targeted therapies have revolutionized cancer treatment. However, in many cases single-agent targeting of mutated oncogenes shows less than optimal results, as both intrinsic and acquired resistance can limit the overall effectiveness of these agent. Efforts to identify secondary therapies have focused on either identifying synthetic lethal targets to enhance therapeutic efficacy and overcome intrinsic resistance or identifying and treating acquired resistance after it occurs. In contrast, studies that assess the extent to which drug combinations can delay the onset of acquired resistance and thereby prolong the initial treatment window are often lacking due to the unavailability of cell culture methods to assess acquired resistance. Here, we describe an *in situ* resistance assay, performed in a 96-well culture format, that allows easy assessment of therapeutic resistance and isolation of therapy-resistant populations.

As a model for assay development, we used osimertinib resistance in *EGFR*- mutated LUAD. Osimertinib is well-established as first-line therapy for patients with EGFR-mutated LUAD^2, 3^, the mechanisms driving acquired resistance are well defined^4-^ _13_, and mechanism-based treatment of acquired resistance is largely unsuccessful since most patients develop tumors harboring multiple parallel resistance mechanisms simultaneously^14^. We found that osimertinib resistance could be reliably modeled in *EGFR*-mutated LUAD cell lines using doses of osimertinib defined by the individual cell lines dose-response curves. Further, osimertinib-resistance populations could be isolated from resistance assays that showed RTK-dependent resistance mechanisms similar to patient populations, but targeting these resistance mechanisms with RTK inhibitors did not reliably re-sensitize cells to osimertinib. In contrast, pan-RTK inhibition using an inhibitor of the proximal RTK signaling intermediate SHP2 synergistically enhanced the efficacy and potency of osimertinib across osimertinib resistant populations. However, even though SHP2 inhibition synergized with osimertinib to kill resistant populations, combined EGFR/SHP2 inhibition produced only ∼70% killing, highlighting the need for combinations that block the development of resistance rather than treating resistant tumors.

A majority of osimertinib resistance is driven by RTK reactivation, thus, pan-RTK inhibition has the potential to circumvent the development of most forms of osimertinib resistance. Indeed, we show that SHP2 inhibition both delayed the onset of osimertinib resistance and limited the overall percentage of populations able to become osimertinib resistant. Downstream of RTKs, the SHP2 phosphatase acts as an adaptor to recruit the RASGEFs SOS1 and SOS2 to receptor complexes and promote RAS activation. SOS1 inhibition limits both intrinsic and acquired resistance to KRAS^G12C^ inhibitors^45^, suggesting that proximal RTK inhibition may be a general strategy to overcome resistance to targeted therapies in lung adenocarcinoma^46^.

Finally, we show our approach to assessing osimertinib resistance *in situ* can be easily adopted to multiple additional RTK/RAS pathway inhibitors with similar objectively defined drug-dosing parameters (Fig. 6). While studies that identify synergistic drug targets to boost therapeutic efficacy to RTK/RAS pathway inhibitors abound, there has been a paucity of assays to systematically assess the extent to which these same combinations can limit acquired resistance. We propose that *in situ* resistance assays, performed over a 6–12-week period, fill this gap in our arsenal of studies assessing combination therapies for cancer treatment. Further, the ease of testing multiple combinations simultaneously, combined with the recent decision by the FDA to no longer requires animal testing for approval of new drugs^47^, make *in situ* resistance assays a cost-effective approach when evaluating therapeutic combinations.

**Fig 6.**
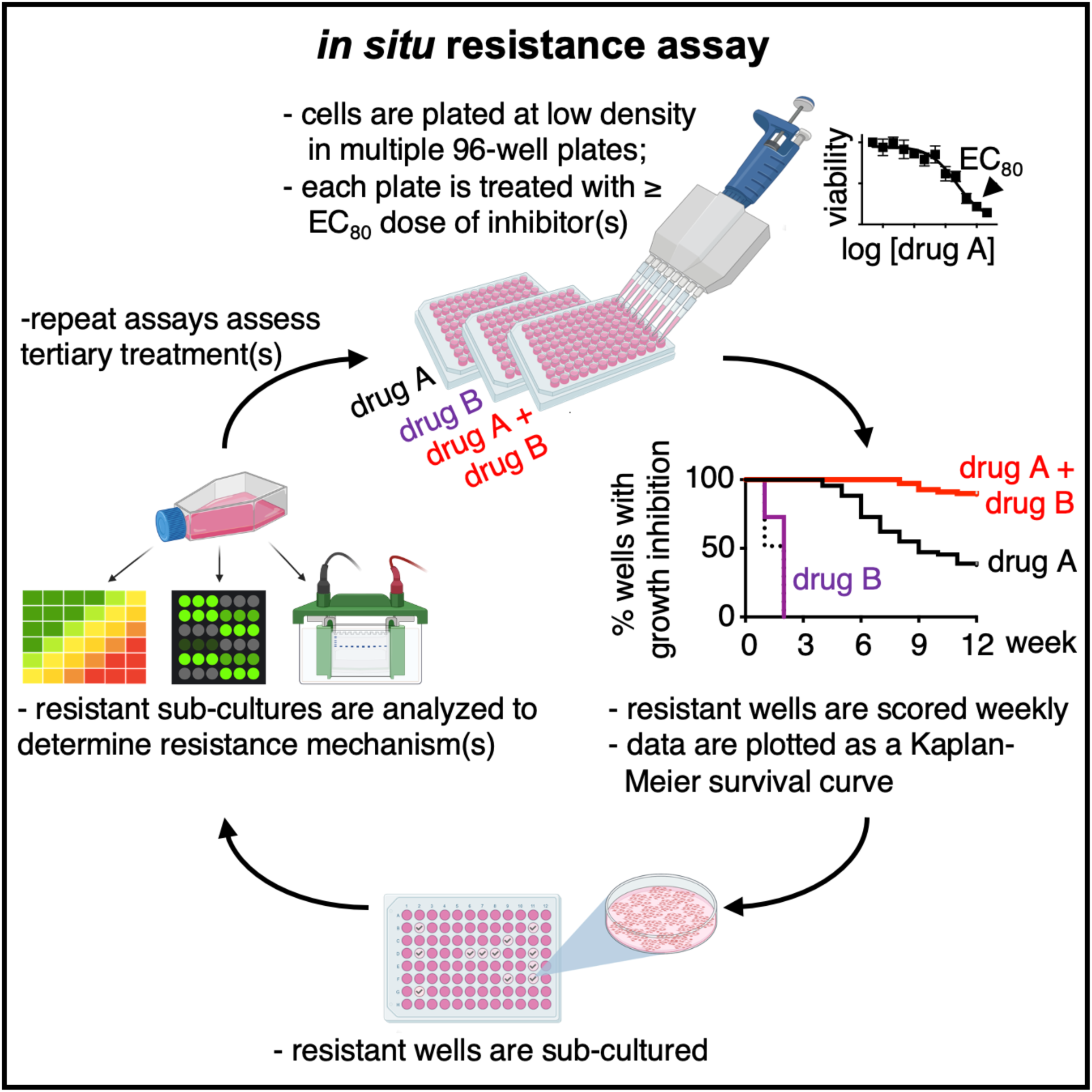
Framework for assessing acquired resistance *in situ*.

## Methods

### Cell culture

Cell lines were cultured at 37°C and 5% CO_2_. HCC827, NCI-H1975, PC9, PC9-TM, H358, H1373, H727, H1838, and H1915 cells were maintained in Roswell Park Memorial Institute medium (RPMI), SMS-CTR cells were maintained in Dulbecco’s Modified Eagles Medium (DMEM), each supplemented with 10% fetal bovine serum and 1% penicillin-streptomycin. Osimertinib-resistant (OR1-7) H1975 cells were isolated from ISRAs treated with 150 nM osimertinib for > 6 weeks. OR1-7 populations were maintained under osimertinib selection during expansion.

### Inhibitor studies

#### Single dose-response studies

For adherent studies, cells were seeded at 500 cells per well in 100 μL in the inner-60 wells of 96-well white-walled culture plates (Perkin Elmer) and allowed to attach for 24 hours prior to drug treatment. For 3D spheroid studies cells were seeded at 500-1,000 cells per well in 100 μL in the inner-60 wells of 96-well ultra-low attachment round bottomed plates (Corning #7007) or Nunc Nucleon Sphera microplates (ThermoFisher # 174929) and allowed to coalesce as spheroids for 24-48 hours prior to drug treatment. Cells were treated with drug for 96 hours prior to assessment of cell viability using CellTiter-Glo® 2.0. Data were analyzed by non-linear regression using Prism 9.

#### Assessment of synergy

Cells were seeded at 200 cells per well in 40 μL in the inner-312 wells of 384-well white-walled culture plates (Perkin Elmer) and allowed to attach for 24-hours prior to drug treatment. To assess the effectiveness of secondary therapies +/- osimertinib, cells were treated with increasing doses of crizotinib, cabozantinib, infigratinib, selpercatinib, linsitinib, SGI-7079, dasatinib, or RMC-4550 +/- 10 nM osimertinib. To assess the extent to which each secondary inhibitor enhanced the effectiveness of osimertinib, cells were treated with increasing doses of osimertinib +/- 100 nM or 300 nM of one of the above inhibitors. To assess drug-drug synergy across a matrix of dose combinations for the osimertinib/RMC-4550 combination, cells were treated with increasing doses of RMC-4550 alone, osimertinib alone, or osimertinib + RMC-4550 in a 6×10 matrix of drug combinations on a similog scale. Cells were treated with drug for 96 hours prior to assessment of cell viability using CellTiter-Glo® 2.0. Values were normalized to DMSO controls and assessment of synergy via Bliss Independence analysis was completed as reported previously ^48^. To deconvolute synergistic synergy versus potency, data were analyzed by Multi-dimensional Synergy of Combinations (MuSyC) Analysis ^33, 34^ using an online tool (https://musyc.lolab.xyz) and are presented as mean +/- 95% confidence interval.

### In situ resistance assays (ISRAs)

Cells were seeded at 250 cells/well in 100 μL in replicate 96-well tissue culture plates and allowed to adhere for 24 hours prior to drug treatment. Plates were then either fed with an additional 100 μL of media or treated with the indicated single dose of targeted inhibitor (osimertinib, sotorasib, adagrasib, trametinib, or tipifarnib) corresponding to ∼ EC_50_ – EC_85_ dose of that drug. For Fig. 3, plates were treated with the indicated drug combinations. The inner 60 wells of each plate were assessed weekly for signs of cell growth using a 4 x objective, with wells reaching >50% confluence scored as resistant. Plates were fed weekly; media was removed from the entire plate in a cell culture hood by quickly inverting the plate into an autoclave bin lined with paper towels to avoid cross-contamination of wells and plates were fed with a multi-channel repeating pipettor. Data are plotted as a Kaplan-Meyer survival curve; significance was assessed by comparing Kaplan-Meyer curves using Prism 9.

#### PhosphoTyrosine Array Analysis

Cells were lysed in RIPA buffer (1% NP-40, 0.1% SDS, 0.1% Na-deoxycholate, 10% glycerol, 0.137 M NaCl, 20 mM Tris pH [8.0], protease (Biotool #B14002) and phosphatase (Biotool #B15002) inhibitor cocktails) for 20 minutes at 4°C and spun at 10,000 RPM for 10 minutes. Clarified lysates were diluted to 1 mg/mL in RIPA buffer prior to assessing for RTK phosphorylation using a Human RTK Phosphorylation Array G1 (RayBiotech, AAH-PRTK-G1-8) per manufacturer’s instructions. Slides were shipped to RayBiotech for data acquisition. For each phosphorylated protein, data were normalized to the average level of phosphorylation observed from two independent isolates of H1975 parental lysate.

## Acknowledgments

This work was supported by funding from the NIH (R01 CA255232 and R21 CA267515 to RLK) and the CDMRP Lung Cancer Research Program (LC180213 to RLK). The funders had no role in the study design, data collection and interpretation, or the decision to submit the work for publication. We thank Dr. Edward Stites (Yale School of Medicine) for helpful discussions during the manuscript’s preparation. The opinions and assertions expressed herein are those of the authors and are not to be construed as reflecting the views of Uniformed Services University of the Health Sciences or the United States Department of Defense. Materials are available upon request from RLK.

## Author Contributions

NES, PLT, and RLK designed the experiments and analyzed the data; NES, PLT and RLK performed most of the experiments; AJL and BRD performed resistance assays; JEH assisted with dose response curves. NES and RLK wrote the manuscript, BRD edited the manuscript.

