## Supplemental Figures S1 - S4 for "In situ modeling of acquired resistance to RTK/RAS pathway targeted therapies"

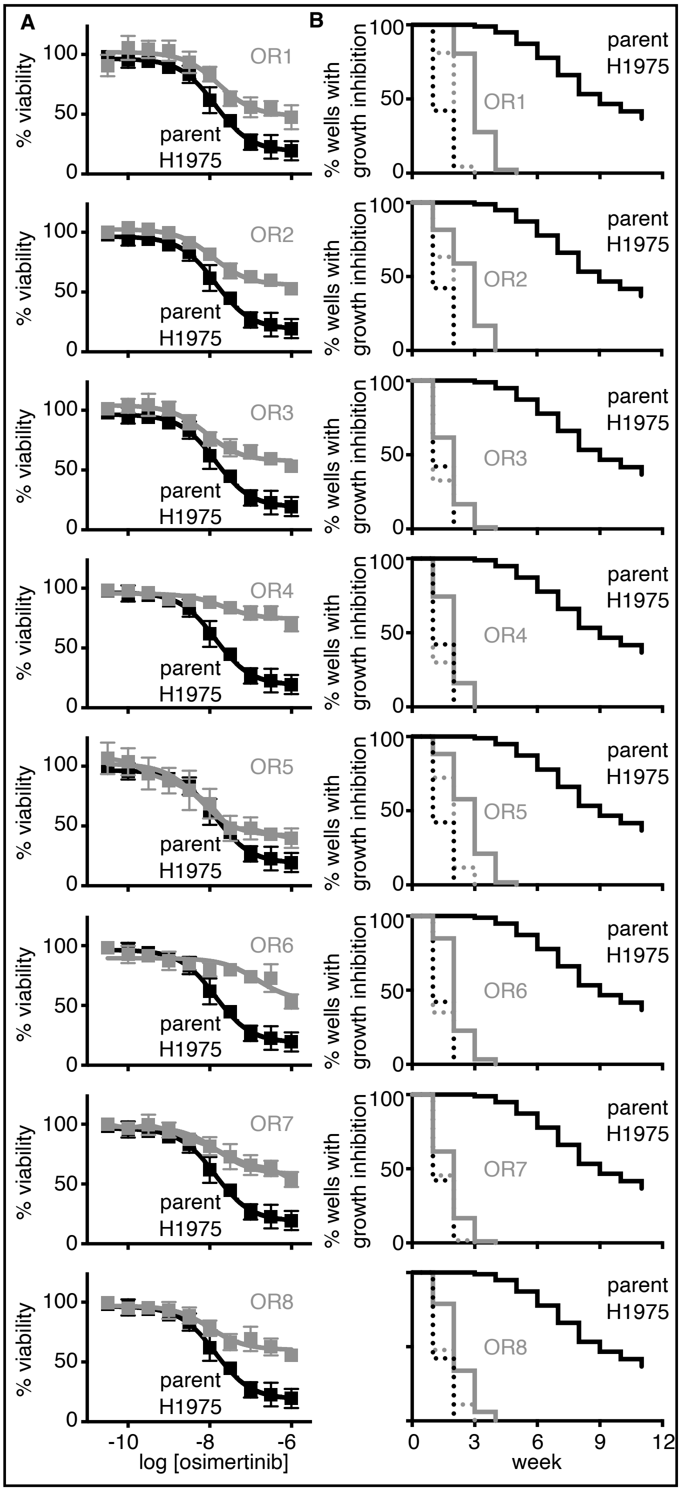


**Fig S1. Isolated osimertinib resistant clones maintain osimertinib resistance upon continued culture.**
**A.** Dose response curves for osimertinib in parental H1975 cells (black) and osimertinib-resistant H1975 populations (grey) isolated from osimertinib resistance assays shown in Fig. 3.
**B.** H1975 parental (black) or osimertinib-resistant populations (grey) were plated at low density (250 cells/well) in replicate 96-well plates each plate was either left untreated (dashed line) or treated with 150 nM osimertinib (solid line). Wells were fed and assessed weekly for outgrowth, wells that were >50% confluent were scored as osimertinib resistant. Data are plotted as a Kaplan-Meyer survival curve.

**A**

osimertinib-resistant H1975 cells

| | H1975 | OR1 | OR2 | OR3 | OR4 | OR5 | OR6 | OR7 | OR8 |
|---|---|---|---|---|---|---|---|---|---|
| Control | 1.0 | 1.0 | 1.0 | 1.0 | 1.0 | 1.0 | 1.0 | 1.0 | 1.0 |
| ABL1 | 1.0 | 1.8 | 1.7 | 1.3 | 1.5 | 1.7 | 1.4 | 2.1 | 1.2 |
| ACK1 | 1.0 | 1.1 | 1.2 | 1.3 | 1.2 | 1.3 | 1.2 | 1.0 | 1.2 |
| ALK | 1.0 | 1.3 | 0.8 | 1.0 | 1.1 | 1.6 | 1.2 | 1.0 | 0.8 |
| AXL | 1.0 | 2.4 | 3.1 | 1.9 | 2.1 | 2.4 | 2.2 | 1.6 | 2.2 |
| BLK | 1.0 | 1.2 | 1.8 | 1.3 | 1.1 | 1.0 | 1.1 | 1.2 | 1.3 |
| BMX | 1.0 | 0.9 | 0.9 | 0.8 | 0.8 | 1.0 | 0.9 | 1.0 | 0.8 |
| BTK | 1.0 | 1.1 | 1.4 | 1.1 | 0.9 | 1.0 | 1.0 | 1.3 | 0.8 |
| CSK | 1.0 | 1.1 | 1.2 | 1.1 | 1.1 | 1.2 | 1.0 | 1.0 | 1.0 |
| TDK | 1.0 | 1.2 | 2.0 | 1.2 | 1.0 | 1.2 | 1.2 | 1.3 | 1.0 |
| EGFR | 1.0 | 1.6 | 1.0 | 1.2 | 1.1 | 1.4 | 0.5 | 0.5 | 1.0 |
| EPHA1 | 1.0 | 1.2 | 2.4 | 1.1 | 1.0 | 1.0 | 1.1 | 2.3 | 1.1 |
| EPHA2 | 1.0 | 1.5 | 1.4 | 1.2 | 1.5 | 1.7 | 1.2 | 1.5 | 0.9 |
| EPHA3 | 1.0 | 1.0 | 1.1 | 0.9 | 0.9 | 1.0 | 1.1 | 1.0 | 0.8 |
| EPHA4 | 1.0 | 0.8 | 1.2 | 1.0 | 1.2 | 1.3 | 1.4 | 1.0 | 1.1 |
| EPHA5 | 1.0 | 1.0 | 0.9 | 0.9 | 0.9 | 0.9 | 0.9 | 0.9 | 1.0 |
| EPHA6 | 1.0 | 0.9 | 0.9 | 0.8 | 1.0 | 0.9 | 1.0 | 1.0 | 0.9 |
| EPHA7 | 1.0 | 0.9 | 0.9 | 0.8 | 0.8 | 0.9 | 0.7 | 1.0 | 0.9 |
| EPHA8 | 1.0 | 1.0 | 1.0 | 0.9 | 0.9 | 1.0 | 1.0 | 0.9 | 0.9 |
| EPHB1 | 1.0 | 1.0 | 1.0 | 0.9 | 0.9 | 1.0 | 0.9 | 0.9 | 0.9 |
| EPHB2 | 1.0 | 1.4 | 2.4 | 1.4 | 1.4 | 1.4 | 1.4 | 1.4 | 1.4 |
| EPHB3 | 1.0 | 2.0 | 1.9 | 1.4 | 1.8 | 2.1 | 2.0 | 1.7 | 1.2 |
| EPHB4 | 1.0 | 1.4 | 1.2 | 1.0 | 1.1 | 2.0 | 1.0 | 1.0 | 0.6 |
| EPHB6 | 1.0 | 1.5 | 1.4 | 0.8 | 0.8 | 1.0 | 0.7 | 1.0 | 1.1 |
| ERBB2 | 1.0 | 1.1 | 1.2 | 1.0 | 1.0 | 1.1 | 1.1 | 1.0 | 1.0 |
| ERBB3 | 1.0 | 1.3 | 2.1 | 0.9 | 0.9 | 1.0 | 1.2 | 1.5 | 1.2 |
| ERBB4 | 1.0 | 1.1 | 1.1 | 0.9 | 0.9 | 1.1 | 0.9 | 1.3 | 0.9 |
| FAK | 1.0 | 1.0 | 0.8 | 0.7 | 1.0 | 1.1 | 0.8 | 0.9 | 0.7 |
| FER | 1.0 | 0.9 | 0.8 | 0.5 | 0.8 | 0.7 | 0.5 | 0.3 | 0.9 |
| FGFR1 | 1.0 | 0.8 | 0.9 | 0.8 | 0.9 | 0.8 | 0.8 | 0.9 | 0.8 |
| FGFR2 | 1.0 | 1.0 | 1.1 | 0.8 | 1.0 | 1.1 | 1.1 | 1.0 | 1.0 |
| FGFR2-a | 1.0 | 2.6 | 1.5 | 2.1 | 2.2 | 3.7 | 2.3 | 1.3 | 1.4 |
| FGR | 1.0 | 1.6 | 1.3 | 1.1 | 1.2 | 1.6 | 1.4 | 1.6 | 0.8 |
| FRK | 1.0 | 1.0 | 1.0 | 0.8 | 0.9 | 1.0 | 0.9 | 0.9 | 0.9 |
| FYN | 1.0 | 1.5 | 2.0 | 1.3 | 0.9 | 1.0 | 1.1 | 1.4 | 1.2 |
| HCK | 1.0 | 1.0 | 1.1 | 0.9 | 0.9 | 1.0 | 1.0 | 1.0 | 1.0 |
| c-MET | 1.0 | 1.5 | 1.4 | 0.9 | 1.1 | 1.2 | 1.0 | 1.0 | 0.9 |
| IGF-1R | 1.0 | 1.6 | 2.2 | 1.1 | 1.2 | 1.0 | 1.3 | 1.6 | 1.3 |
| Insulin-R | 1.0 | 1.0 | 1.0 | 0.8 | 0.9 | 1.0 | 0.8 | 0.9 | 0.8 |
| ITK | 1.0 | 1.0 | 1.0 | 0.9 | 1.1 | 1.1 | 1.1 | 0.9 | 0.9 |
| JAK1 | 1.0 | 1.3 | 0.9 | 1.0 | 0.9 | 2.0 | 1.3 | 1.0 | 0.8 |
| JAK2 | 1.0 | 1.2 | 1.2 | 1.0 | 1.0 | 0.8 | 0.7 | 1.0 | 0.9 |
| JAK3 | 1.0 | 1.1 | 0.7 | 0.8 | 0.9 | 1.3 | 1.0 | 0.8 | 0.6 |
| LCK | 1.0 | 1.3 | 1.5 | 1.0 | 1.1 | 1.5 | 1.5 | 1.4 | 1.3 |
| LTK | 1.0 | 1.6 | 1.9 | 1.4 | 1.5 | 1.6 | 1.4 | 1.7 | 1.2 |
| LYN | 1.0 | 1.0 | 1.1 | 1.1 | 0.9 | 1.2 | 1.0 | 1.1 | 0.9 |
| MATK | 1.0 | 0.9 | 1.0 | 1.0 | 0.9 | 1.1 | 1.0 | 0.9 | 0.9 |
| M-CSFR | 1.0 | 1.0 | 1.3 | 0.7 | 0.9 | 1.0 | 0.8 | 1.1 | 0.9 |
| MUSK | 1.0 | 1.0 | 1.0 | 0.9 | 0.8 | 1.2 | 1.0 | 1.0 | 1.1 |
| NGFR | 1.0 | 1.2 | 1.2 | 1.2 | 1.3 | 1.7 | 1.3 | 1.2 | 1.1 |
| PDGFR-a | 1.0 | 1.1 | 1.3 | 1.0 | 1.0 | 1.2 | 1.1 | 1.0 | 0.9 |
| PDGFR-b | 1.0 | 0.9 | 1.0 | 0.8 | 0.8 | 0.9 | 0.7 | 1.0 | 0.8 |
| PYK2 | 1.0 | 1.1 | 1.6 | 0.7 | 1.0 | 1.4 | 1.0 | 1.5 | 0.7 |
| RET | 1.0 | 1.8 | 1.1 | 1.2 | 1.3 | 2.3 | 1.8 | 1.3 | 1.0 |
| ROR1 | 1.0 | 1.5 | 2.0 | 1.3 | 1.3 | 1.4 | 1.2 | 1.1 | 1.1 |
| ROR2 | 1.0 | 1.8 | 1.5 | 1.4 | 1.3 | 1.2 | 1.2 | 1.0 | 1.0 |
| ROS | 1.0 | 1.2 | 1.1 | 0.9 | 1.0 | 1.1 | 0.9 | 1.1 | 1.0 |
| RYK | 1.0 | 1.5 | 1.4 | 0.8 | 1.2 | 1.6 | 1.3 | 1.7 | 0.8 |
| c-KIT | 1.0 | 1.8 | 1.6 | 1.1 | 0.7 | 1.2 | 1.2 | 1.6 | 1.5 |
| SRMS | 1.0 | 1.0 | 1.0 | 1.1 | 1.1 | 1.2 | 0.9 | 0.9 | 0.9 |
| SYK | 1.0 | 1.0 | 1.2 | 0.8 | 0.9 | 1.1 | 0.9 | 1.0 | 0.9 |
| TEK | 1.0 | 2.0 | 2.1 | 0.9 | 1.5 | 0.8 | 1.0 | 1.0 | 1.3 |
| TIE1 | 1.0 | 1.5 | 1.9 | 1.1 | 1.4 | 1.5 | 1.3 | 1.9 | 1.1 |
| TIE2 | 1.0 | 1.4 | 1.6 | 0.9 | 0.9 | 1.0 | 1.0 | 1.7 | 0.8 |
| TNK1 | 1.0 | 1.0 | 0.9 | 0.9 | 0.8 | 1.2 | 1.0 | 1.2 | 0.9 |
| TRKB | 1.0 | 1.2 | 1.0 | 0.9 | 0.9 | 1.5 | 1.3 | 1.0 | 0.8 |
| TXK | 1.0 | 1.4 | 1.2 | 1.1 | 1.1 | 1.8 | 1.3 | 1.1 | 0.8 |
| TYK2 | 1.0 | 1.0 | 1.0 | 0.9 | 1.0 | 1.0 | 1.0 | 0.9 | 0.9 |
| TYRO10 | 1.0 | 1.1 | 1.4 | 0.8 | 1.0 | 1.2 | 1.1 | 1.8 | 0.9 |
| VEGFR2 | 1.0 | 1.0 | 1.7 | 0.8 | 1.0 | 0.9 | 1.0 | 1.3 | 1.0 |
| VEGFR3 | 1.0 | 1.3 | 1.7 | 0.9 | 1.1 | 1.3 | 1.3 | 1.5 | 0.9 |
| ZAP70 | 1.0 | 0.9 | 0.8 | 0.7 | 1.0 | 1.6 | 1.0 | 1.0 | 0.7 |

**B**

| inhibitor | **target**, other targets |
|---|---|
| crizotinib | **c-MET**, **ROS1**, ALK |
| cabozantinib | **VEGFR2, c-MET, RET, c-KIT, TIE2, AXL** |
| infigratinib | **FGFR1/2/3** |
| selpercatinib | **RET**, VEGFR1/3, FGFR1/2/3 |
| linsitinib | **IGF-1R**, Insulin Receptor |
| SGI-7079 | **AXL**, SYK, JAK2, TRKA/B, PDGFRβ, RET |
| dasatinib | **ABL**, **SRC/SFKs**, c-KIT |
| RMC-4550 | **SHP2** |

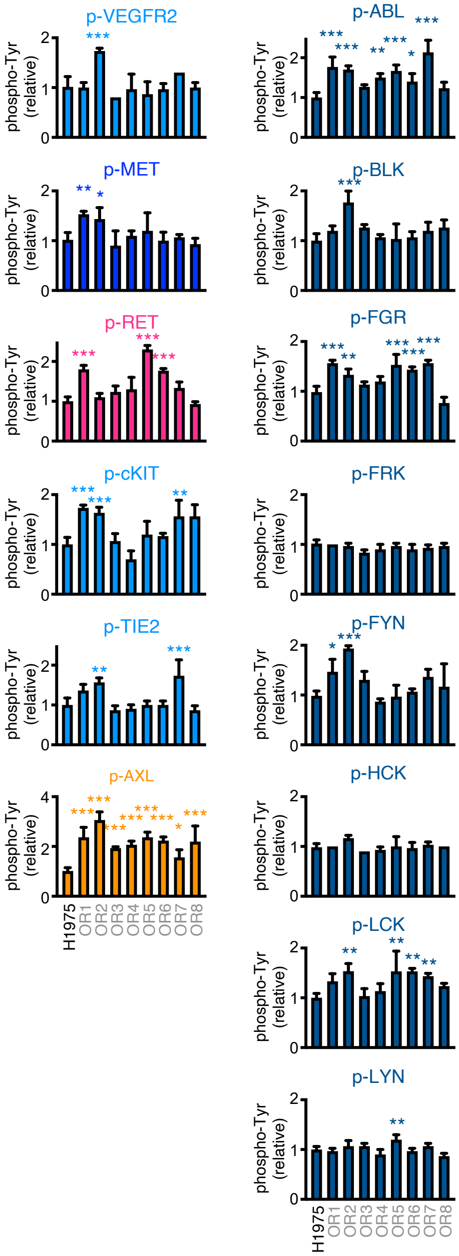


**Fig S2 (related to Fig. 4). Osimertinib resistant populations show activated RTKs (related to Fig 4).**

**A.** Whole cell lysates from parental H1975 cells (black) and osimertinib-resistant H1975 populations from Fig. 3 (grey) were assessed for RTK phosphorylation using a Human RTK phosphorylation antibody array (RayBiotech). Data were normalized to the average of two independent isolates of parental H1975 lysates. Red numbers indicate increased tyrosine phosphorylation of the indicated kinase over parental controls. Colored arrows indicate drug targets for inhibitors in **B** that were tested in combination with osimertinib in Fig. 4; large arrows indicate the primary drug target whereas small arrows indicate the multiple secondary targets for a given color coded inhibitor.

**B.** RTK and RTK/RAS pathway inhibitors assessed for synergy in combination with osimertinib in Fig. 4. The protein targets for each inhibitor are indicated, **bold** indicated the primary drug target for each inhibitor.

**C-D.** Quantitation of tyrosine phosphorylation for the multiple targets of cabozantinib (**C**) and dasatinib (**D**). Primary target phosphorylation is repeated from Fig. 4. Cabozantinib targets that are the primary target for another drug listed in B are color coded as a target of that drug and are repeated from Fig. 4 for completeness.

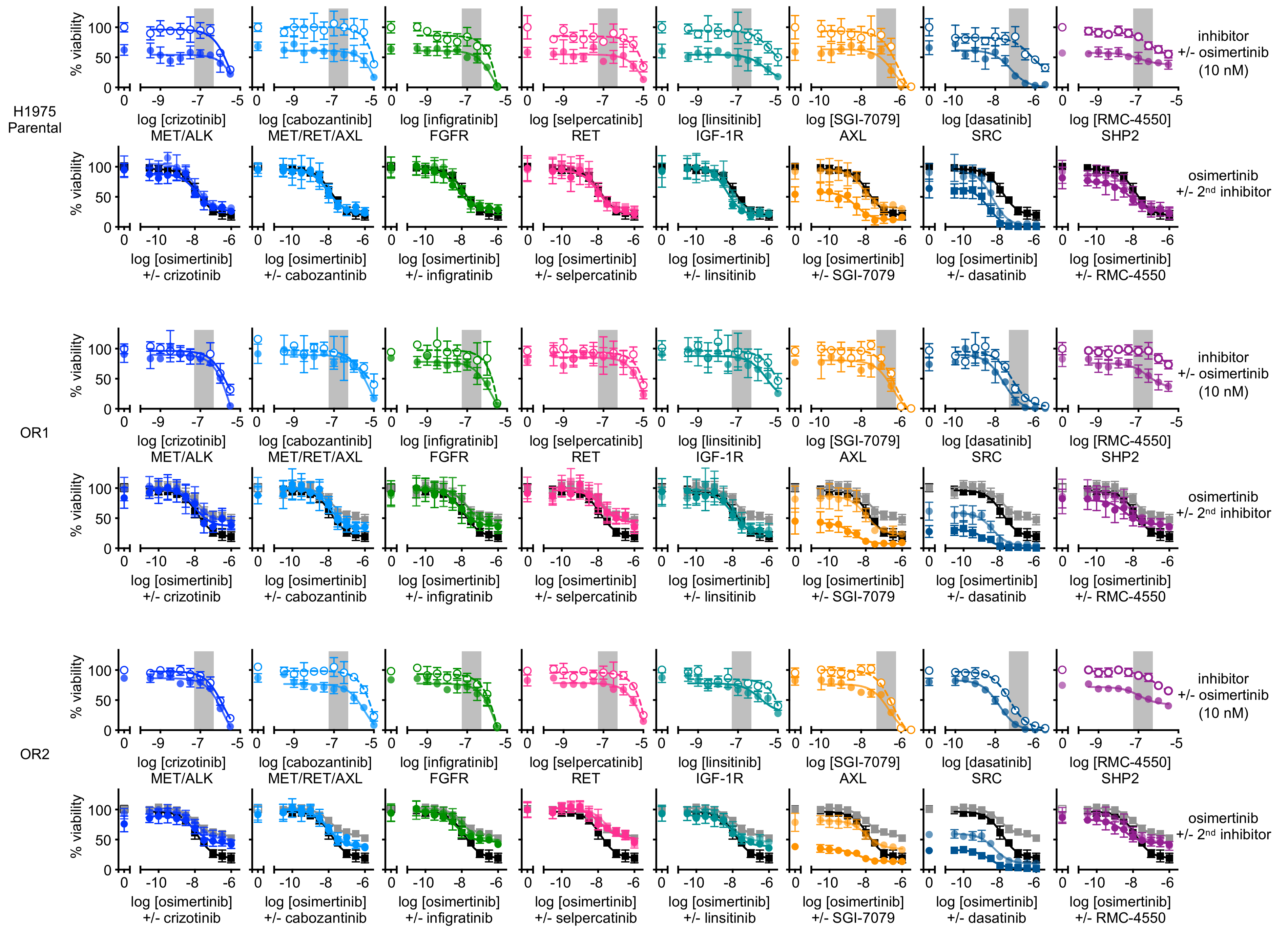
H1975 Parental
OR1
OR2
% viability
log [crizotinib] MET/ALK
log [cabozantinib] MET/RET/AXL
log [infigratinib] FGFR
log [selpercatinib] RET
log [linsitinib] IGF-1R
log [SGI-7079] AXL
log [dasatinib] SRC
log [RMC-4550] SHP2
inhibitor +/- osimertinib (10 nM)
log [osimertinib] +/- crizotinib
log [osimertinib] +/- cabozantinib
log [osimertinib] +/- infigratinib
log [osimertinib] +/- selpercatinib
log [osimertinib] +/- linsitinib
log [osimertinib] +/- SGI-7079
log [osimertinib] +/- dasatinib
log [osimertinib] +/- RMC-4550
osimertinib +/- 2nd inhibitor

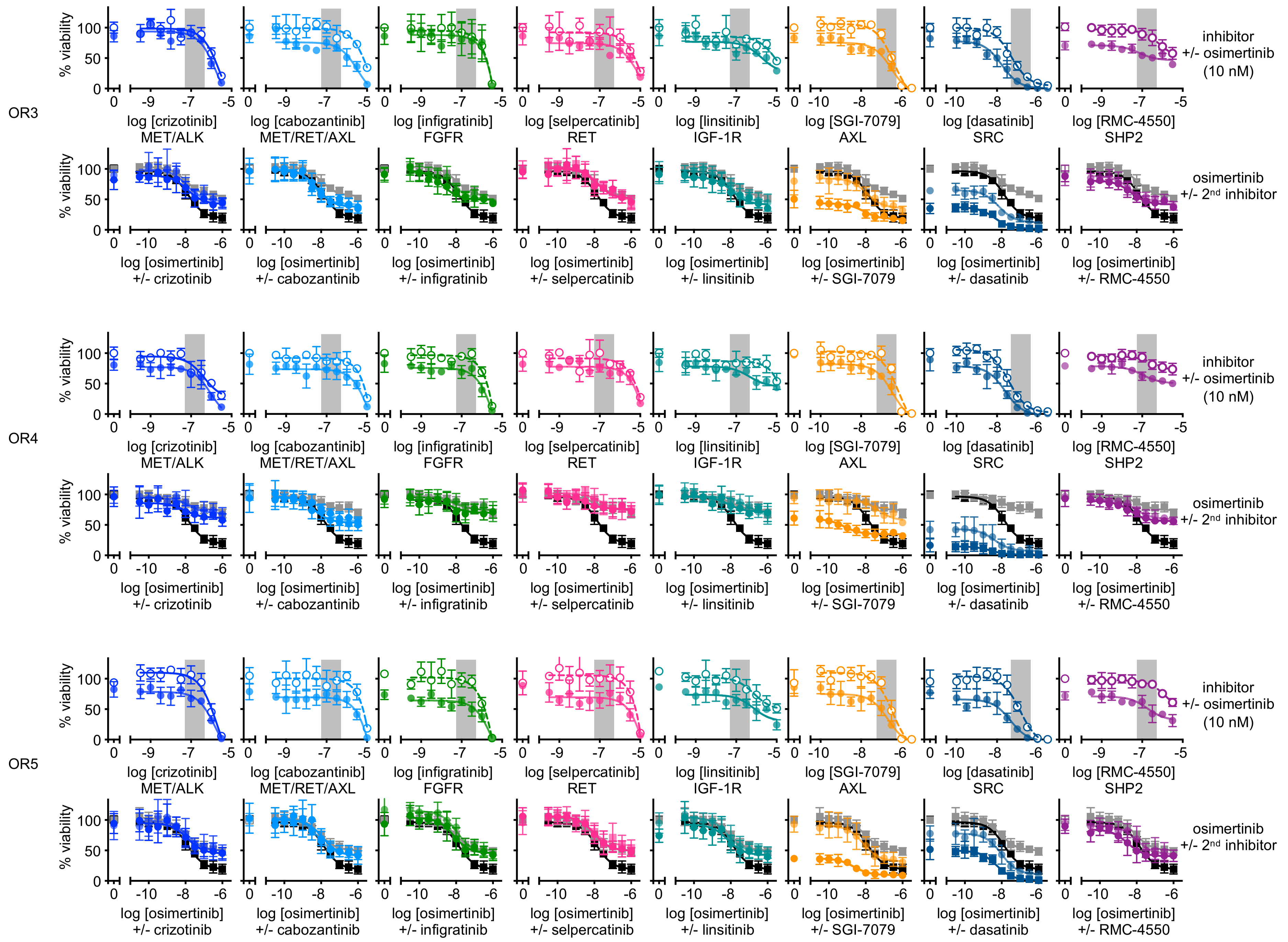
OR3
OR4
OR5
% viability
inhibitor +/- osimertinib (10 nM)
osimertinib +/- 2nd inhibitor
log [crizotinib] MET/ALK
log [cabozantinib] MET/RET/AXL
log [infigratinib] FGFR
log [selpercatinib] RET
log [linsitinib] IGF-1R
log [SGI-7079] AXL
log [dasatinib] SRC
log [RMC-4550] SHP2
log [osimertinib] +/- crizotinib
log [osimertinib] +/- cabozantinib
log [osimertinib] +/- infigratinib
log [osimertinib] +/- selpercatinib
log [osimertinib] +/- linsitinib
log [osimertinib] +/- SGI-7079
log [osimertinib] +/- dasatinib
log [osimertinib] +/- RMC-4550

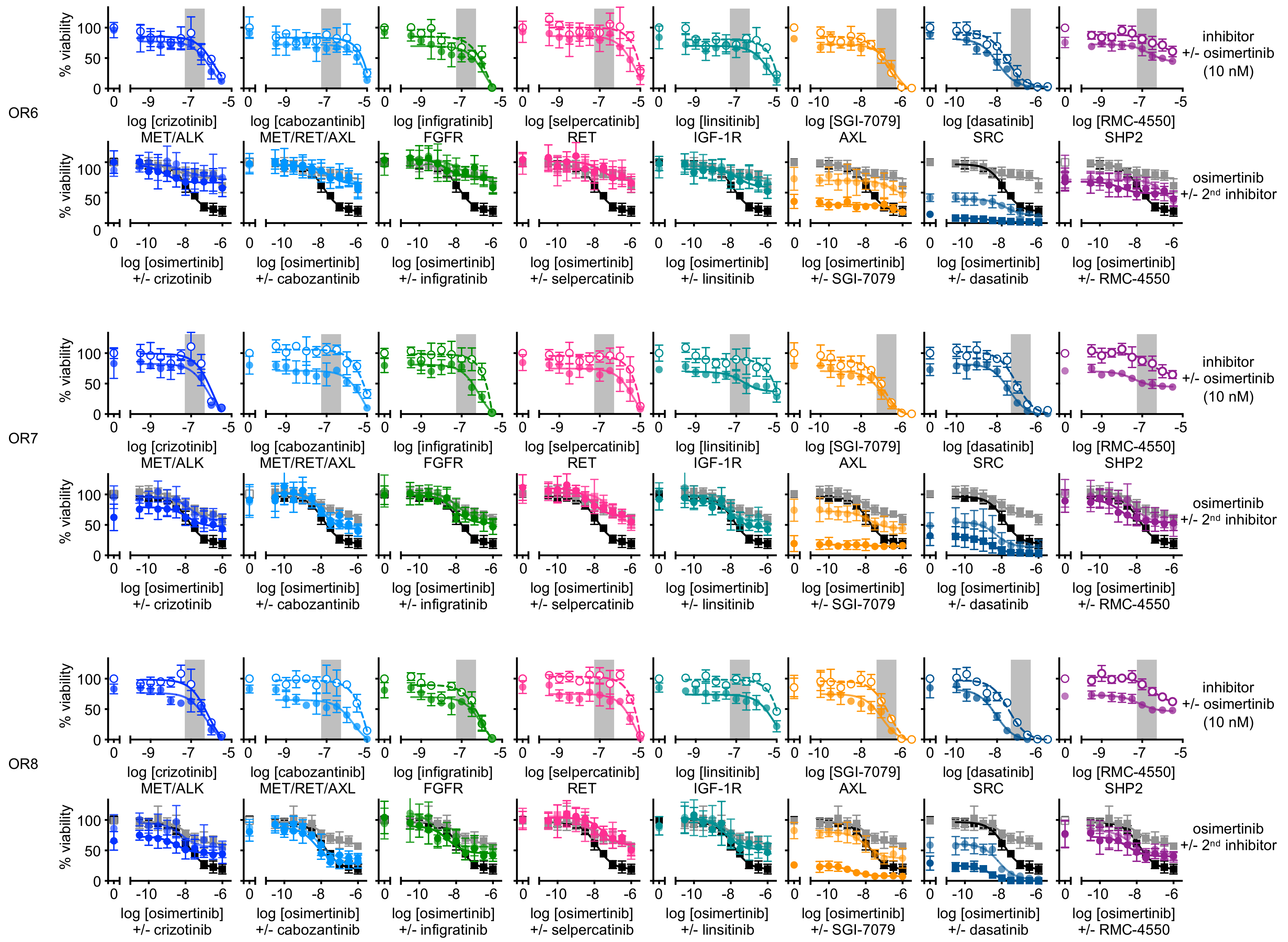
OR6
OR7
OR8
% viability
log [crizotinib]
MET/ALK
log [cabozantinib]
MET/RET/AXL
log [infigratinib]
FGFR
log [selpercatinib]
RET
log [linsitinib]
IGF-1R
log [SGI-7079]
AXL
log [dasatinib]
SRC
log [RMC-4550]
SHP2
log [osimertinib]
+/- crizotinib
+/- cabozantinib
+/- infigratinib
+/- selpercatinib
+/- linsitinib
+/- SGI-7079
+/- dasatinib
+/- RMC-4550
inhibitor
+/- osimertinib
(10 nM)
osimertinib
+/- 2nd inhibitor

**Fig S3 (related to Fig. 4). Isolated osimertinib resistant populations show differential responses to combining osimertinib with RTK inhibitors versus inhibitors of downstream (SFK, SHP2) signaling.**
Dose response curves of parental and osimertinib-resistant H1975 populations were treated either increasing doses of a second RTK/RAS pathway –targeted inhibitor +/- 10 nM osimertinib (top) or with increasing doses of osimertinib +/- the second RTK/RAS pathway-targeted inhibitor at 100 or 300 nM (bottom) for four days. The greyed box on the top indicates the targeted inhibitor doses used in combination with the osimertinib dose response curves on the bottom plots. Cell viability was assessed using CellTitre glo. Synergistic efficacy and potency calculations for each drug combination based on MuSyC analysis are shown in Fig. 2B.

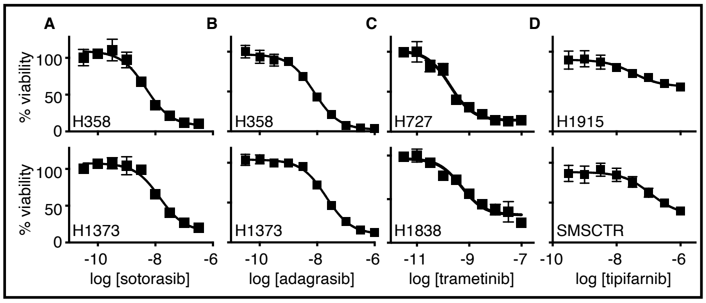


**Fig S4 (related to Fig. 5). Dose response curves for cells treated with sotorasib, adagrasib, trametinib, or tipifarnib.** **A-D**. Dose response curves for *KRAS*$^{G12C}$-mutated H358 and H1373 cells (A, B), *KRAS*$^{G12C}$-mutated H727 and NF1-LOF H1838 cells (C), or HRAS-mutated H1915 and SMSCTR cells treated increasing doses of sotorasib (A), adagrasib (B), trametinib (C), or tipifarnib (D). Sotorasib, adagrasib, and trametinib sensitivities were assessed in 3D cultured spheroids; tipifarnib sensitivity was assessed in 2D adherent culture. $EC_{50}$ – $EC_{90}$ values from these dose response curves are shown in Fig. 5E.
